# Protocol for Non-viral HDR-based CRISPR/Cas9 platform for small custom editing in primary T cells

**DOI:** 10.64898/2026.02.01.703084

**Authors:** Katariina Mamia, Anniken Solveig Matheson Sollano, Shiva Dahal-Koirala, Emma Haapaniemi

## Abstract

CRISPR/Cas9 enables precision gene editing via HDR for mutation correction and disease modelling. This protocol describes an 8-day non-viral HDR workflow for editing primary patient and healthy donor T cells, including reagent design, editing, on-target detection, and flow cytometry. The protocol was developed under research-grade conditions but supports scaling up and the transition to preclinical and clinical GMP workflows. For complete details on the use and execution of this protocol, please refer to Mamia et al.[1].

**Before you begin:** CRISPR/Cas9 gene editing is a promising tool to correct pathogenic variants for autologous cell therapies, targeting monogenic diseases such as inborn errors of immunity (IEI). Furthermore, it can be used as a tool for disease modelling to study normal and pathological variations of the immune system. Here we present a detailed protocol for an efficient and customizable T cell single nucleotide variant (SNV) correction platform based on homology-directed repair (HDR). The protocol details every step of the process, which starts with custom CRISPR/Cas9 reagent design of guide-RNAs (gRNAs) and repair templates for editing a novel target with no previously published reagents. Furthermore, we describe the strategy of reagent design to assess on-target HDR editing using droplet digital PCR (ddPCR). Next, we detail the T cell platform itself, and present effective strategies to stimulate PBMCs *ex vivo* to promote CD4+ and CD8+ T cell activation and proliferation, which we have validated in 32 unique IEI patients. Next, we present the workflow of gene editing T cells using nucleofection and CRISPR ribonucleoprotein (RNP) complexes for efficient editing that preserves high cell viabilities and up to 80% HDR. Finally, we present a flow cytometry panel that assesses the immune cells present at the end of the platform, including characterization of memory and effector T cell populations and status of T cell exhaustion.

**Innovation:** In the study, we present a detailed protocol of performing highly efficient, non-viral and HDR-based precision editing in patient and healthy control T cells. The developed platform enables custom editing, such as correction of small pathogenic variants, with one workflow that we demonstrate to achieve up to 80% efficiency in multiple genomic loci and donors.

**Institutional permissions:** The study was conducted in accordance with the principles of the Helsinki Declaration and approved by the Helsinki University Central Hospital Ethics Committee, and the Regional Committee for Medical and Health Research Ethics South-East Norway. All participants have signed written informed consent.

## 1.1 Target sequence identification and visualization

### Timing: [variable]

This step outlines the initial phases of precision editing a novel locus, where either a wild-type sequence is used to design editing tools for healthy controls, or a patient-specific sequence is used for mutation correction or another type of edit. For a novel target that does not have previously published CRISPR/Cas9 reagents, the protocol starts here. If previously published CRISPR/Cas9 reagents are available, proceed to Step 2.

1. Mutation information is provided using different versions of the human reference genome assembly. We recommend using the GRCh38/hg38 assembly, along with the genomic coordinates (i.e. chromosome number: precise position on the chromosome).
2. Alternatively, the mutation information is based on the transcript. This nomenclature includes details about the reference transcript (e.g., NM_007181), the exon where the mutation is located (e.g., exon 23), the coding DNA coordinate (e.g., c.1778), and a description of the nucleotide change (e.g., T>G).
  a. **Note:** A single gene can have multiple transcripts, and therefore different coding DNA coordinates. Always make sure to use the correct transcript ID.
3. Go to Ensembl.org and search for the gene of interest.
4. Copy the assembly ID (written next to the gene name).
  1. Go to SnapGene, select “Import” and “Ensembl sequences” and import gene by pasting the copied ID from Ensembl.
    a. **Note:** Alternative gene visualization tools can be used from this step forward. Please follow instructions of the specific tool.
  2. If the patient-specific sequence is available, align it in SnapGene to verify that the sequence is matching to find the exact position of the mutation:
    a. Alignment: Ctrl+F or “Edit”→”Find”→”Find DNA Sequence”.
  3. If patient-specific sequence is not available, add it manually:
    a. Go to https://genome.ucsc.edu/ and select “Genome Browser”.
    b. Select GRCh38/hg38 in human assembly drop-down menu and the genomic position in “Position/Search Term”.
    c. The mutated nucleotide is highlighted in yellow. Use zoom out function for better visualization.
    d. In the menu on the bottom of the page, enable NCBI RefSeq: <u>pack</u> (or according to the preference) to obtain information on exon/intron, codon, amino acid, and position.
  4. Reduce the length of the target sequence to +/- 400 bp from the mutation/target site and create two files for CRISPR reagent design:
    a. File that shows the sequence after editing
    b. File that shows the sequence before editing (e.g. WT/MUT sequence)

## 1.2 CRISPR gRNA design and visualization

### Timing: [variable]

This step outlines the design of CRISPR/Cas9 gRNAs for *in vitro* validation. We recommend testing 2-5 gRNAs per target site in primary T cells to identify gRNAs with high HDR efficiency (Fig. 1).

1. Open SnapGene file containing the patient/wild-type sequence intended for editing.
  a. **Note:** It is important to have the correct sequence for gRNA design as efficient gRNAs for pathogenic variants tend to overlap the mutations [1] .
2. Identify possible PAM sites within the target site by searching for “GG” to highlight available PAM sites (5’- NGG-3’) for SpCas9.
  a. **Note:** For homozygous recessive mutations, we recommend screening all available gRNAs within 10-20 bp from the mutation site to limit the number of gRNAs to validate *in vitro* [1].
  b. **Note:** For heterozygous mutations, we strongly recommend designing gRNAs that overlap the mutation site in the gRNA seed region (i.e. the first 10 nt upstream of the PAM site) to reduce risk of cutting the healthy allele.
3. gRNA sequence is the 20 nt sequence directly upstream of the PAM site.
  a. **Note:** gRNAs with high likelihood of off-target sites should be eliminated from further testing, using gRNA off-target prediction tools, such as CHOPCHOP [2, 3].
  b. **Note:** Both forward and reverse gRNAs can be used, we recommend matching the ssODN orientation with the gRNA orientation to achieve high on-target editing.
4. Mark Cas9 cut site on the gRNA design (3 bp upstream from the PAM site).
5. Order gRNAs with sgRNA formulation (ready-to-use) or as a crRNA that is annealed 1:1 with a tracrRNA by following the vendor’s instructions.

**Figure 1.**
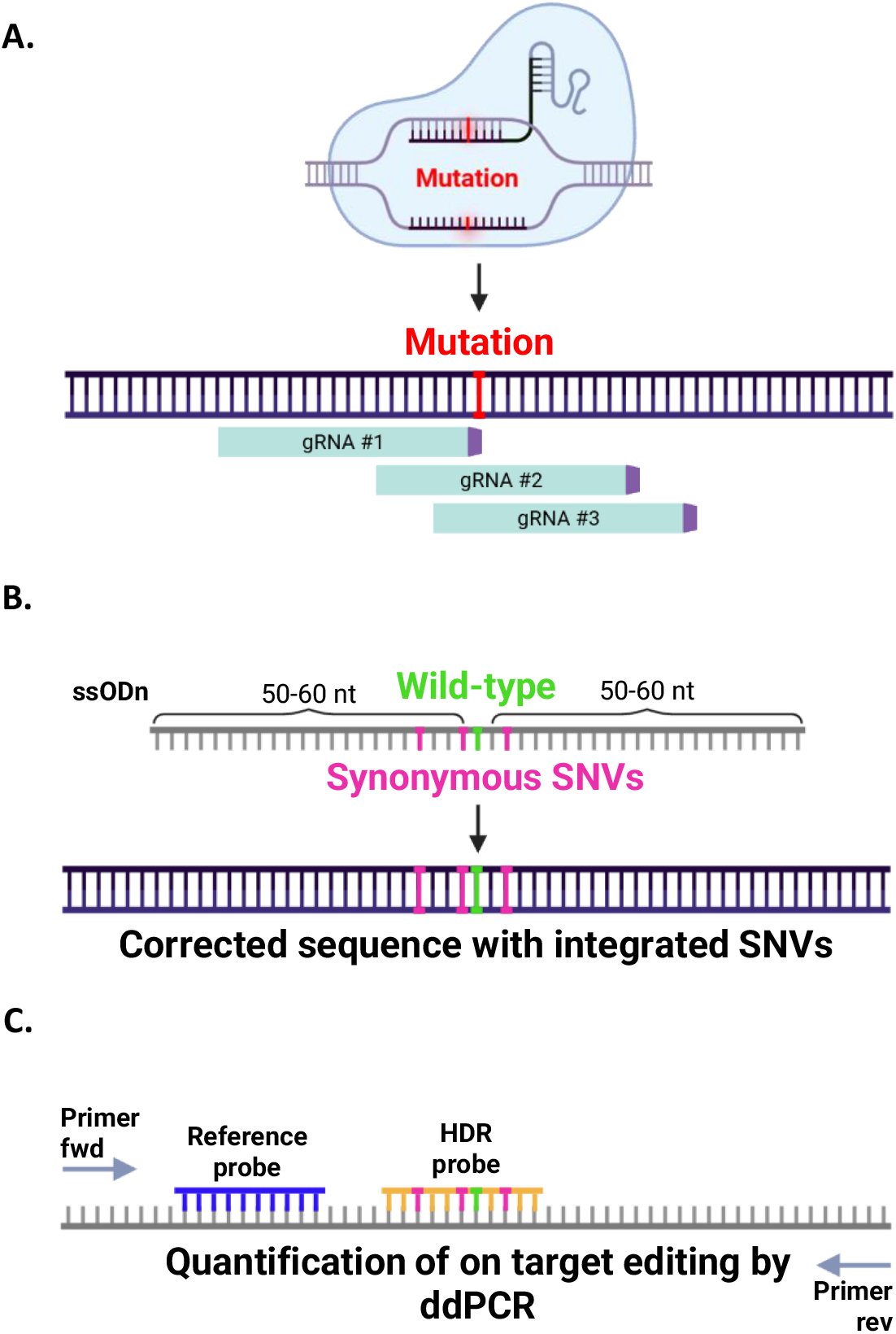
CRISPR/Cas9 reagent design and on-target assessment. **(A)** Schematic representation of the gRNA screening strategy, where multiple gRNAs (green) per target site are designed based on the available PAM sites (purple) within the 100 bp ssODN area. Mutation site is marked with red. Schematic representation of the repair strategy, where 100-120 bp ssODNs with +/-60 bp homology arms from the mutation site (red) are used. ssODN design includes correction of the mutation (green) and 2-3 silent SNVs (pink), enabling identical editing strategy in patients and healthy controls and HDR detection by ddPCR. **(C)** Schematic representation of the ddPCR assay design for HDR detection, where forward and reverse primers amplify a ∼250 bp region centering the editing site. Reference probe (blue) binds to unedited sequence outside of the repair template and HDR probe (orange) binds to the edited sequence, which allows for precise quantification of HDR in edited T cells.

## 1.3 CRISPR ssODN design and visualization

### Timing: [variable]

This step outlines the ssODN design steps for HDR-based editing (e.g. mutation correction) at the target site. In order to prevent Cas9 from re-cutting the edited strand [4, 5], we recommend implementing 2-3 synonymous SNVs into the ssODN design in addition to mutation correction (Fig. 1) [1] . When implementing silent SNVs to the repair template design, we recommend adding them at the PAM site and or the gRNA seed region (i.e. the first 10 nt upstream of the PAM site). This strategy also helps to increase accuracy of ddPCR for HDR detection (see Step 8 below).

1. Design 100-120 nt long ssODN for the target site using the following guidelines:
  a. The ssODN should center the mutation/target site with 50-60 nt homology arms on both sides surrounding the mutation.
  b. The additional 2-3 SNVs should be synonymous, where the nucleotide change does not change the amino acid sequence.
  c. The SNVs should not be located at a highly conserved nucleotide position.
    i. **Note:** Conserved regions can be visualized with UCSC genome browser (https://genome.ucsc.edu/).
  d. SNVs should not be in the intron region.
  e. SNVs should affect splicing.
    i. **Note:** Predictions can be performed using SpliceAI (https://spliceailookup.broadinstitute.org/#), CADD scoring (https://cadd.gs.washington.edu/score) and Human Splicing Finder Pro (https://genomnis.com/hsf).
  f. The SNVs and the mutation/target site should not be more than 10-20 bp apart.
2. Add 2X phoshorothioate (PT) modifications on the 3’ end of the ssODN to improve HDR by preventing the repair template from 3’ endonuclease degradation [1] .

## 1.4 ddPCR reagent design to assess on-target HDR editing

### Timing: [variable]

This step outlines the design of the reagents for assessing on-target editing by ddPCR (Fig. 1). Alternatively, commercially designed kits (e.g. Bio-Rad GED HDR Assay) can be used. We note that ddPCR assay is reliable for HDR detection [1] but does not as accurately capture NHEJ. Alternatively, we recommend assessing on-target editing with NGS-based methods, such as amplicon sequencing, which we have described previously [1, 6].

1. ddPCR primer design:
  a. Use a 300 bp sequence length with +/- 150 bp arms from the mutation site for primer design with a chosen software (such as Primerblast, Primer3 etc.)
  b. Set parameters for primer design: length at 100-300 bp, Tm 54-58 °C.
    i. **Note:** primers should amplify a region long enough to include the repair template entirely as well as the ddPCR probes (reference, HDR and optionally NHEJ)
  c. Choose 2-3 designs and test primers on a 2% agarose gel to confirm primer specificity.
    i. **Note:** To test primers, we recommend validating them using DNA from the target cell type (T cells).
    ii. **Note:** We recommend testing different PCR annealing temperatures (e.g. +/- 5 °C of the primer Tm) to find optimal PCR conditions (Step 8).
    iii. **Note:** consider using a restriction enzyme to reduce unspecific amplification. 2-5 U of restriction enzyme is recommended per sample and is added in the ddPCR mastermix with other reagents (Step 8).
2. General ddPCR probe design guidelines:
  a. Primer sequence cannot overlap the probe sequence.
  b. Probe Tm should be 3-10 °C higher than primer Tm.
  c. GC content should be 30-80%.
  d. Probe length should be <30 nt, ideally 20 nt. Shorter probes are better at discriminating between single nucleotides.
  e. The 5’ end of the probe should not start with a G (this quenches the fluorescence)
3. Reference ddPCR probe design:
  a. Reference probe should be located outside of the target editing site and ideally outside the repair template.
    i. **Note:** External reference probe within another locus can alternatively be used in the assay. However, we did not find differences in ddPCR readouts when comparing internal and external reference probes [1].
  b. Reference probe should be ordered with 5’ HEX and 3’BHQ1 fluorochromes.
    i. **Note:** Other quenching fluorochromes can be used. Follow vendor guidelines.
4. HDR ddPCR probe design:
  a. HDR probe should center the mutation/target site and contain the sequence of the edit, not the unedited sequence.
  b. HDR probe should be ordered with 5’ FAM and 3’BHQ1 fluorochromes.
    i. **Note:** Other quenching fluorochromes can be used. Follow vendor guidelines.
5. Optional: NHEJ ddPCR probe design:
  a. NHEJ probe should center the Cas9 cut site and contain the sequence of the unedited target.
  b. NHEJ probe should be ordered with 5’ FAM and 3’BHQ1 fluorochromes.
    i. **Note:** Other quenching fluorochromes can be used. Follow vendor guidelines.

### 2. Patient/healthy control PBMC isolation from whole blood using density grade centrifugation

#### Timing: [3 h]

This step describes the process of PBMC isolation from whole blood samples collected in sodium heparin tubes. The process results in a clean PBMC population that does not contain plasma, platelets or red blood cells.

1. Pipette 15 mL Lymphoprep per LeucoSep tube and briefly spin down (short spin, 30 s) to move the Lymphoprep below the filter line.
2. Pool 10-20 mL of whole blood from sodium heparin blood collection tubes into a 50 mL Falcon tube.
  a. **Note:** If blood sampling volume is higher, divide blood into multiple 50 mL Falcon tubes.
3. Rinse blood collection tubes with RT PBS and add into the previously prepared 50mL tube(s).
4. Further fill the 50 mL tube containing the blood up to 40 mL with RT PBS to dilute the blood, mix by pipetting.
5. Pipette diluted blood (40 mL per tube) into the LeucoSep tube by gently dispensing the liquid onto the tube wall.
  a. **Note:** Be careful with pipetting (use low dispensing speed for) in order to prevent blood from falling below the filter.
6. Centrifuge LeucoSep tubes for 20 min at 800 g with brakes off (1) and acceleration off (1).
  a. **Note:** The centrifugation takes approximately 30 min to finish.
7. After centrifugation, remove some (∼15-20 mL) of the plasma layer on top with a pipette.
  a. **Note:** This helps to access the PBMC layer better in the next step.
8. Harvest the enriched PBMC cell fraction below the plasma layer by aspirating the cells with a 1000 μL pipette tip into clean 50 mL Falcon tubes pre-filled with ∼20 mL RT PBS.
  a. **Note:** We recommend that contents of one LeukoSep tube goes into one clean 50 mL tube for optimal washing in the next step.
  b. **Note:** Do not aspirate Lymphoprep below the PBMC layer as it can be toxic for the cells.
9. Fill each tube up to 50 mL with RT PBS and mix gently by pipetting.
10. Centrifuge for 10 min at 100 g with brakes on (9) acceleration on (9).
11. Discard supernatant by aspirating with a pipette.
  a. **Note:** PBMC pellet is large but easily dislodged. Do not discard supernatant by pouring.
12. Resuspend cell pellet gently first in 1000 μL RT PBS and then fill up to 50 mL. Mix by pipetting.
  a. See Troubleshooting Problem 1 if PBMCs are aggregated and difficult to dislodge by pipetting at this stage.
  b. See Troubleshooting Problem 2 if a high amount of red blood cells are present at this stage.
13. Centrifuge for 5 min at 300 g with brakes on (9) acceleration on (9).
14. Discard supernatant like previously described and resuspend pellets first in 1000 μL RT PBS. Pool pellets from tubes together and then fill up to 50 mL PBS.
15. Take an aliquot for counting to record cell count and viability.
  a. **Note:** If PBMC yield is high, cells can be diluted before counting to ensure accurate measurement.
16. Centrifuge cells for 5 min at 300 g with brakes on (9) acceleration on (9).
17. Discard supernatant and proceed to cryopreservation or cell culture.

### 3. PBMC cryopreservation

#### Timing: [30 min]

This step describes PBMC cryopreservation for long-term storage at -150°C prior to thawing the cells for editing experiments. We recommend cryopreserving a minimum of 5 million cells/vial (patients/small sampling volumes) and a maximum of 30 million cells/vial (healthy controls/large sampling volumes) for optimal cell recovery.

1. Based on the cell count, prepare to freeze 5-30 million cells per cryovial. Freezing volume per vial is 1 mL unless cell number is <5 million, which is when reduced freezing volumes can be used (0.5 mL).
2. Label cryovials with appropriate information, including donor/patient number, date and number of cells per vial.
3. Resuspend cell pellet from Step 2.17 first in 1 mL of 4 °C CryoStor freezing medium using a 1000 μL pipette, then fill up to a desired volume.
  a. **Note:** DMSO is toxic for the cells. When freezing cells, perform steps quickly while remaining gentle with pipetting.
  b. **Note:** Pipetting with a 1000 μL tip at the beginning is recommended to ensure cells are in single-cell suspension in the freezing medium for optimal recovery.
4. Add 1 mL cell suspension per cryovial, close caps tightly and place vials into CoolCell container.
5. Move container immediately to -80 °C for o/n storage.
  a. **Note:** Do not store cells at -80 °C for more than 1-4 days to ensure optimal recovery.
  b. **Note:** Do not stack CoolCell containers on top of each other at -80 °C to ensure equal freezing between the containers.
6. Transfer cryovials to -150 °C for permanent storage.

### 4 PBMC thawing and cytokine stimulation (day 1)

#### Timing: [1 h]

This step describes the thawing and stimulation of cryopreserved PBMCs. This step is not necessary if PBMCs are cultured directly after isolation described in Step 2. PBMCs are stimulated with a cytokine cocktail, which we have shown previously to promote CD4+ and CD8+ T cell activation and proliferation (Fig. 2) [1].

1. Transfer cells from -150 °C onto dry ice.
2. Pipette 10 mL of RT ImmunoCult T cell expansion medium into a 15 mL Falcon tube per each cryovial to be thawed.
3. Thaw cells in a 37 °C water bath for approximately 2 min until only a pea-sized frozen part is left.
  a. **Note:** do not immerse cryovials fully in water to prevent water from getting into the cap as this could result in contamination.
4. Spray cryovial with 70% ethanol, wipe, and transfer into sterile laminar hood.
5. Gently pipette 1000 μL RT ImmunoCult T cell expansion medium into the vial and pour vial contents into the 15ml Falcon pre-filled with 10 mL thawing medium.
  a. **Note:** Do not pipette cells during this step. When pouring cells, make sure vial contents flows directly into the thawing medium and not the wall of the tube.
6. Add 1000 μL thawing medium into cryovial to wash out remaining cells and pool with the previous.
7. Centrifuge cells at 300 g for 5 min.
8. Discard supernatant, resuspend pellet first in 1mL with a 1000 μL tip with pre-warmed Immunocult T cell expansion medium supplemented with anti 15μL/mL CD3/CD28 or 15μL/mL CD3/CD28/CD2 and IL-2 (120 U/mL), IL-17 (3 ng/μL) and IL-15 (3 ng/μL). From here onwards, this medium is referred to as ‘T cell stimulation medium’. Then add up to 1-10 mL depending on the size of the pellet.
  a. **Note:** cells should be at a minimum of 1 million cells/mL at the time of counting to avoid unnecessary centrifugation step.
9. Mix gently by pipetting. Take an aliquot for counting.
10. Record cell viability and amount of living cells/mL
11. Dilute cells into 1 million cells/mL with pre-warmed T cell stimulation medium.
  a. **Note:** We have compared two T cell stimulation cocktails (CD3/CD28 or CD3/CD28/CD2 in healthy controls and 30 IEI patients (Fig. 2).
12. Pipette cell suspension into an appropriate cell culture plate/flask and incubate cells at 37 °C/5 % CO_2_ for three days without disturbing them.
  a. **Note:** We advise to use tissue-culture treated cell culture plates/flasks to promote beneficial cell-to-cell communication between T cells and macrophages during early stages of PBMC stimulation.
  b. **Note:** The choice of cell an appropriate cell culture flask depends on the cell count. We recommend following the guidelines below prevent drying of cells while ensuring proper gas exchange in the flask:
    i. < 1 million: One well in 12-well plate
    ii. < 2 million: One well in 6-well plate
    iii. 2-4 million: T-25 flask standing upright
    iv. 4-6.5 million: T-25 flask lying down
    v. 6.5-10 million: T-75 flask standing upright
    vi. 10-15 million: T-75 flask lying down
    vii. 15-30 million: T-175 flask standing upright
    viii. >40 million: T-175 flask lying down

**Figure 2.**
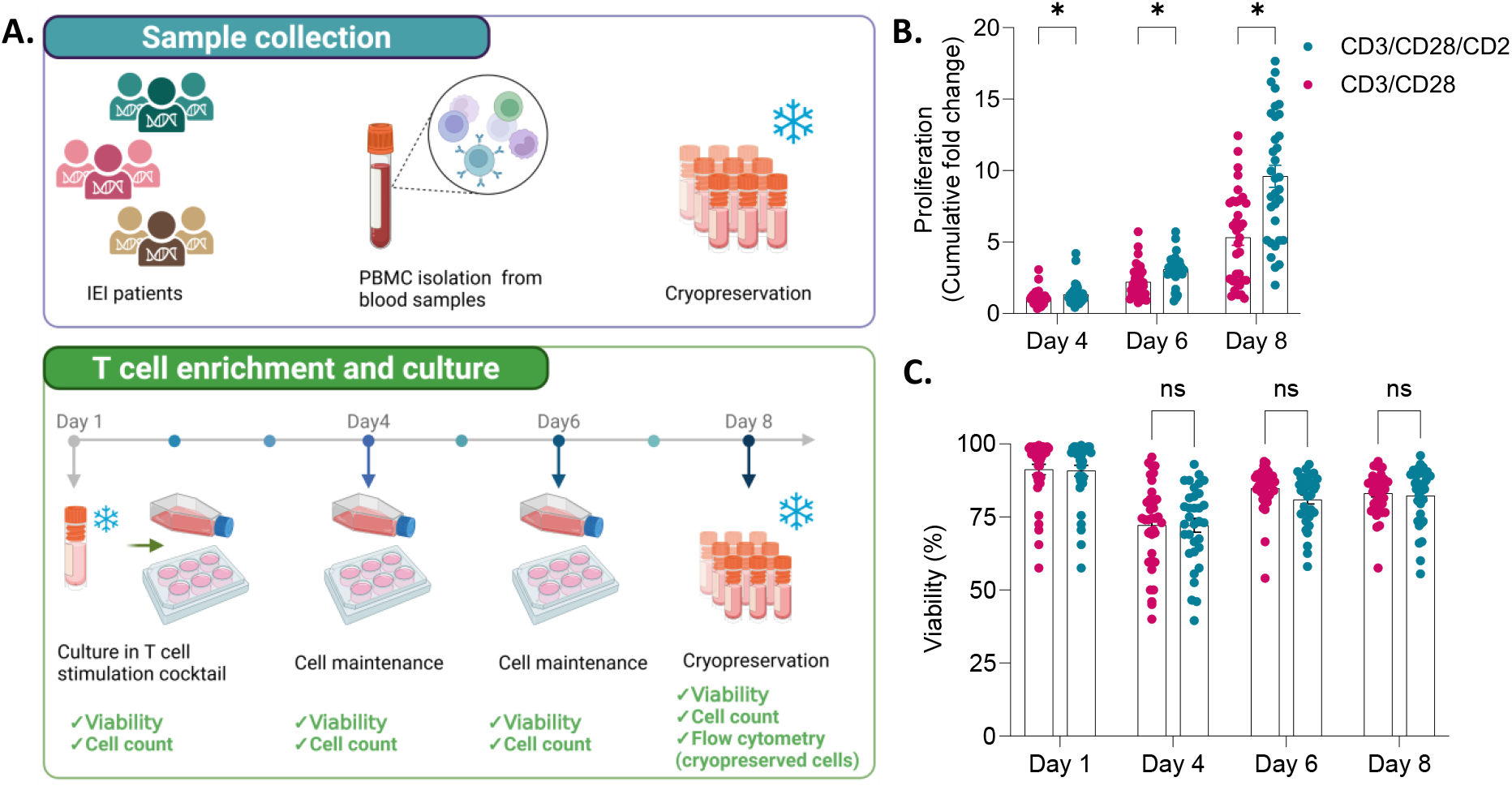
Overview of the T cell platform. (**A**) Schematic of the eight-day T cell enrichment and culture workflow. In brief, PBMCs isolated from blood are cryopreserved, then thawed and cultured with IL-2, IL-7, IL-15, and CD3/CD28 or CD3/CD28/CD2 stimulation. After three days, medium is refreshed with interleukins on days 4 and 6. Cells are cryopreserved on day 8 for flow cytometry. Viability and cell counts are recorded on days 1, 4, 6, and 8. (**B**) Viability and **(C)** cumulative fold expansion relative to day 1 of the cells derived from 33 IEI patients. Data are presented as mean ± SEM. Statistical analyses were performed in GraphPad Prism using the Wilcoxon matched-pairs signed rank test (nonparametric, paired design). P values were adjusted for multiple comparisons using the Holm–Šidák method, with a significance threshold of α = 0.05.

### 5. PBMC dilution (day 4)

#### Timing: [15 min]

This step describes the dilution of the previously stimulated PBMCs. If there are enough cells on day 4, this step is optional, and one can proceed directly to nucleofection. However, when performing large experiments and or working with little starting material, we recommend performing this step prior to nucleofection to expand cells further.

1. Dilute cells at 0.25-0.5 million cells/mL or 1:1 in volume on day 4 by adding pre-warmed T cell stimulation medium without CD3/CD28 or CD3/CD28/CD2.
  a. **Note:** Keep cells in the same flask during this step as they are partially adhered at this stage and expand more thoroughly if cell culture vessel is not changed.
  b. **Note:** Cell counts (especially for patients) at this stage are not reliable due to partially adhered T cells.
  c. **Note:** Cell dilution factor depends on the individual and should be assessed case-by-case. We note that patient cells expand more slowly compared to healthy controls, while healthy controls can be diluted to 0.25 million cells/mL without issues.
  d. d. See Troubleshooting Problem 3 if cell viability is low on Day 4-8.
2. Incubate cells at 37 °C/5 % CO_2_ for two days and repeat dilution on day 6 and 8 if necessary.
  a. **Note:** It is ideal to nucleofect cells between day 4-8 to achieve high precision editing outcomes. After day 8, precise editing levels via HDR decrease rapidly, as we have previously shown [1].
  b. See Troubleshooting Problem 4 if there are not enough cells on day 4 of the protocol

### 6. T cell nucleofection (day 4-8)

#### Timing: [1-3 h depending on the size of the experiment]

This step describes the nucleofection of previously stimulated PBMCs. We recommend performing the nucleofection between day 4-8 of the PBMC stimulation protocol to ensure high HDR levels in the samples [1]. Edited cells can be collected 4 days after nucleofection or expanded further.

1. Pre-program the Lonza 4D electroporation unit for 16-well strip or 96-well plate depending on the size of the experimental setup.
2. Prepare cell culture plates by adding Immunocult T cell expansion medium supplemented with IL-2 (250 U/mL) into the wells. From here onwards, this medium is referred to as ‘T cell recovery medium’. Incubate at 37 °C/5 % CO_2_ while proceeding with the protocol.
  a. **Note:** Cell culture plates and working volumes depend on the number of cells per nucleofected sample. As low as 0.1 million cells per sample can be nucleofected but due to the fragility of patient cells, we recommend opting for 0.5-1 million cells per nucleofected sample.
    i. 0.1-0.25 million cells: 96-well plate with a final volume of 200 uL
    ii. 0.5-0.7 million cells: 48-well plate with a final volume of 400 uL
    iii. 0.8-1 million cells: 24-well plate with a final volume of 500 uL
3. Prepare CRISPR RNPs for nucleofection:
  a. Mix 100 pmol sgRNA or 1:1 annealed crRNA-tracrRNA and 61 pmol Cas9 nuclease per reaction by pipetting or vortexing gently.
    i. **Note:** If multiple different experimental conditions are tested, we recommend preparing the individual RNP reactions in a 96-well plate and sealing the plate with foil.
    ii. **Note:** RNPs can be made for several replicas in one reaction (i.e. in mL tubes) if experimental design allows.
  b. Briefly spin the plate/tube down and incubate at 37 °C for 15 min
  c. Transfer reaction on ice until 100 pmol repair template/reaction is added.
    i. **Note:** repair template should be added when cells are ready for nucleofection to prevent RNP degradation.
4. Prepare cells for nucleofection:
  a. Harvest cells from cell culture flasks and transfer into a 50 mL Falcon tube
    i. **Note:** Additional PBS washing in the flask can be performed to harvest as many cells as possible.
  b. Centrifuge cells at 300 g for 5 min.
  c. Discard supernatant by pipetting and resuspend cell pellet in 10-50 mL PBS depending on the number of cells.
  d. Mix and take an aliquot for counting
  e. Count the volume of cell suspension needed to obtain the desired number of cells for nucleofection (0.1-1 million cells per nucleofected sample).
  f. Transfer desired volume into a new 15 or 50 mL tube and centrifuge at 300 g for 5 min.
    i. **Note:** It is advisable to add the repair template to the RNP mix at this stage.
  g. Discard supernatant by pipetting and resuspend cell pellet in P3 nucleofection buffer with 20 μL buffer per nucleofected sample
    i. **Note:** Depending the nucleofection buffer (we recommend Lonza Primary cell P3 nucleofection buffer for primary T cells), it is advisable to work fast as some buffers can be toxic for primary cells.
    ii. **Note:** It is important that cells are in single cell suspension in nucleofection buffer to ensure successful nucleofection and editing levels.
5. Mix cells with RNPs:
  a. Add 20 μL of cell suspension per sample into the previously prepared RNPs.
    i. **Note:** If multiple different experimental conditions are tested in the experiment, we recommend adding the cells into the RNPs on a 96-well plate.
    ii. **Note:** Several replicas can be made in one reaction (1.5 mL tubes) if the experimental design allows.
  b. Mix gently but thoroughly to obtain a homogenous single-cell suspension.
6. Perform nucleofection:
  a. Using a multichannel pipette (RNPs in a 96-well plate) or an electronic pipette (RNPs in 1.5 mL tubes), transfer 20-25 μL cell suspension per electroporation cuvette well.
    i. **Note:** Cell volume can be adjusted according to the volume of RNPs (RNP volume is usually 3-5 μL per sample depending on the reagent concentrations) but should not exceed 30 μL.
    ii. **Note:** Pipette carefully when transferring cell-RNP suspension into the nucleofection wells as bubbles may disturb the procedure and affect cell viability and editing outcomes.
  b. Nucleofect selected wells using pulse code ‘EO-115’ and solution ‘Primary Cell P3’.
    i. **Note:** Different pulse codes and solutions may be used for different experiments. This setup works for previously stimulated healthy donor and patient T cells.
  c. Immediately after nucleofection, pipette 85 μL pre-warmed T cell recovery medium into each well.
    i. **Note:** Do not pipette cells up and down at this point as they are fragile.
  d. Incubate cells at 37 °C/5 % CO_2_ for 15 min.
7. Transfer nucleofected cells into cell culture plates:
  a. After incubation, add 50 μL of T cell recovery medium into each well
  b. Transfer cells into previously prepared cell culture plates with a multichannel pipette.
    i. **Note:** Do not pipette cells up and down at this point as they are fragile.
  c. Gently move the plates in a ‘criss-cross’ movement to distribute cells evenly without pipetting.
  d. Incubate cells at 37 °C/5 % CO_2_ for 24h.

### 7. T cell maintenance after nucleofection (day 5-8)

#### Timing: [30 min]

This step describes the maintenance protocol for nucleofected T cells. We recommend performing the first dilution/splitting step 24h after nucleofection and the second step 72h after nucleofection. We recommend collecting cells 4 days after nucleofection (day 8 of the platform) as at this point HDR editing levels have stabilized [1] unless the edited cells have a growth advantage/disadvantage and require a timepoint experiment to determine the optimal time for sample collection.

1. 24h after nucleofection, dilute/split cells by adding pre-warmed T cell recovery medium onto the cells.
  a. **Note:** Dilution/splitting ratio depends on the donor and the well size. Healthy controls can typically be split at 1:2 to 1:3 at this stage as the cells tend to grow fast. Depending on the patient, cells can be diluted (i.e. add medium) or split at 1:1 to 1:2.
  b. **Note:** When splitting cells, make sure to dislodge T cell colonies into single-cell suspension by pipetting gently. When working with very fragile donor, keep cells at colonies at this stage to avoid shearing stress on the cells.
  c. **Note:** Optimal cell density after dilution/splitting at this stage is 0.4-1 million cells/mL depending on the donor. Cells density before the next splitting/dilution (72h after nucleofection) should not exceed 2-2.5 million cells/mL to ensure high cell viability.
  d. **Note:** Keep in mind the recommended working volumes in a given well and transfer cells into a larger plate when needed:
    i. 96-well plate: 100-200 μL
    ii. 48-well plate: 200-400 μL
    iii. 24-well plate: 500-1000 μL
    iv. 12-well plate: 1000-2000 μL
2. 72h after nucleofection, repeat cell dilution/splitting while maintaining appropriate T cell density.
  a. **Note:** At this stage, T cell colonies are typically large and dislodging them by gentle pipetting is recommended.
3. 96h after nucleofection, harvest cells for downstream analyses or cryopreservation.
  a. **Note:** Cells can be further expanded by continuing to culture them in T cell recovery medium, which is added every second day.
  b. **Note:** If cells are expanded, we recommend to re-stimulate T cells on day 10-12 of the platform with full T cell stimulation medium. T cells are then seeded at 1 million cells/mL as described in Step 4. After re-stimulation, cells should be diluted with T cell stimulation medium without CD3/CD28 every 2-3 days at 0.25-0.4 million cells/mL depending on the donor/patient.

### 8. On-target editing assessment by ddPCR (day 8 or as preferred) Timing: [8 h]

This step describes the process of sample preparation for ddPCR to detect on-target editing in the samples. We note that the assay is more reliable when in addition to mutation correction/target, 2-3 additional synonymous SNVs have been integrated into the repair template design. This ensures that the HDR probe can discriminate between an unedited and edited sequence, which can be more challenging with only a single nucleotide correction. Alternatively, commercially designed kits (e.g. Bio-Rad GED HDR Assay) can, which have been optimized for single nucleotide correction detection.

1. Extract gDNA from cells as according to kit instructions.
  a. **Note:** we recommend 0.5-1 million cells/sample to achieve high gDNA yield.
  b. **Note:** gDNA samples can be stored at -20°C indefinitely until running the ddPCR assay.
2. Measure gDNA concentration using Qubit or other sensitive DNA concentration measurement kit.
3. Normalize gDNA concentration across samples to 8 ng/μL with nuclease-free ddH_2_O.
  a. **Note:** If DNA stock concentration is <8 ng/uL, it is possible to use lower dilutions for the assay at as low as 1 ng/uL as long as all samples have been normalized to the same concentration.
4. Pipette 8 μL of DNA dilution per well on a 96-well semi-skirted ddPCR plate.
  a. **Note:** ddPCR DNA dilutions can be stored at -20 °C until running the assay.
5. Thaw ddPCR primers, FAM and HEX probes and ddPCR supermix (see design of custom ddPCR reagents in Step 1.4) in room temperature.
  a. **Note:** ddPCR probes are light-sensitive and have a longer shelf-life when they are protected from light.
6. Prepare ddPCR master mix for HDR/NHEJ assay in a DNAse/RNAse free 1.5 mL tube:
  a. Calculate the amount of reagents needed as according to Table 1.
  b. Add nuclease-free ddH_2_O, ddPCR primers, reference probe, HDR/NHEJ probe and mix by vortexing.
  c. Vortex ddPCR supermix vigorously to solubilize any precipitated crystals.
  d. Add ddPCR supermix to the reagents and mix by vortexing.
7. Add 12 μL ddPCR mastermix per sample into 96-well semi-skirted ddPCR plate with the diluted gDNA.
  a. **Note:** If there are empty wells in the column, fill them up with 20 μL of water.
8. Seal the plate, mix by vortexing. Briefly spin down.
9. Generate droplets using Automated Droplet Generator as according to manufacturer’s instructions.
10. Place piercable foil on droplets and perform heat sealing.
11. Run PCR according to Table 1.
  a. **Note:** the program takes about 3.5 h to finish, finished plate can be stored at 4 °C o/n prior to reading the plate.
12. Read droplets in Droplet Reader.
13. Analyze data according to experimental design using QX Manager.
  a. See Troubleshooting Problem 5 if HDR editing is low in the samples (0-10% HDR)
  b. See Troubleshooting Problem 6 if NHEJ editing is low in knockout experiments (i.e. when repair template is not used).

**Table 1.**
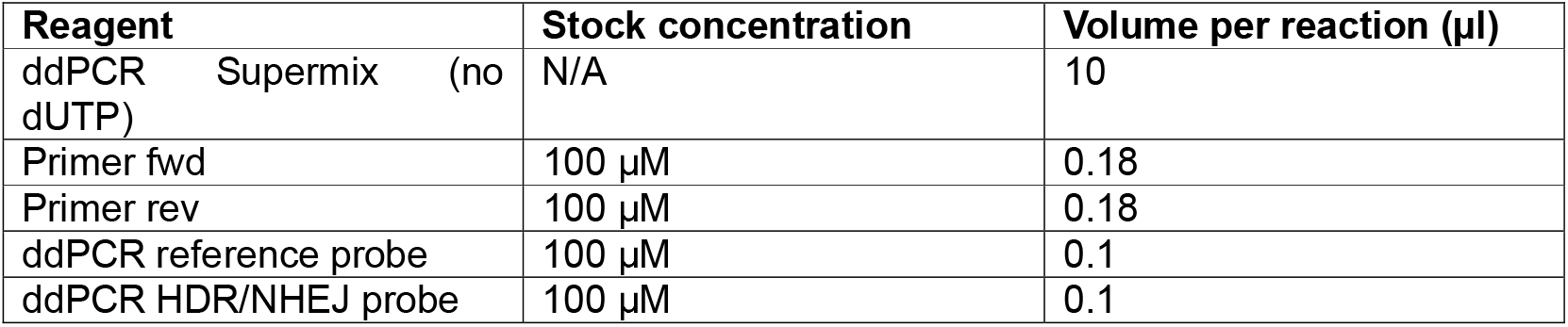
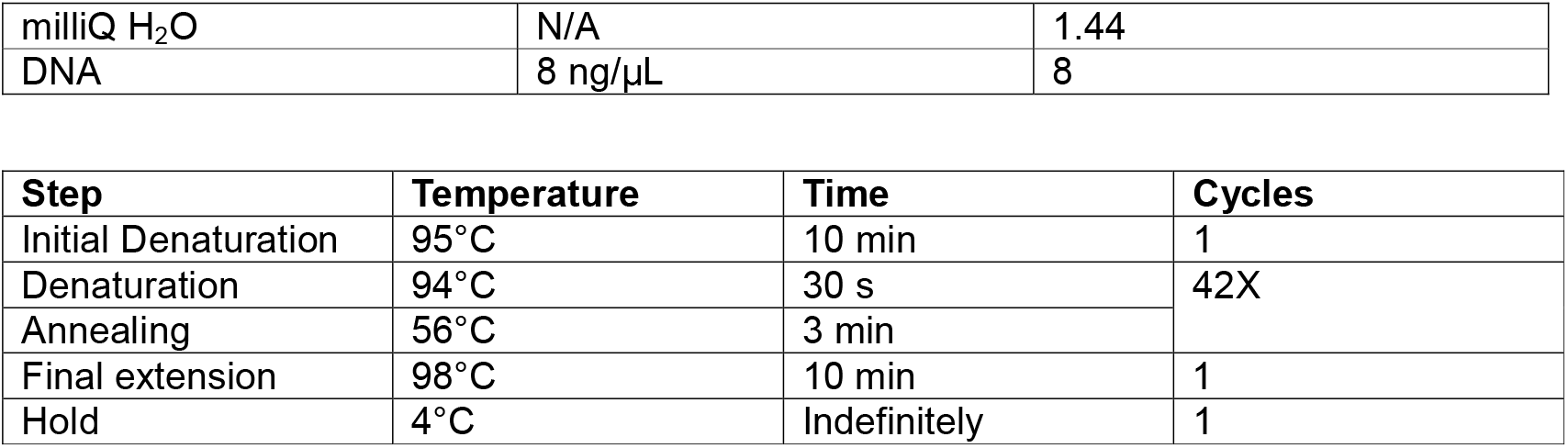
ddPCR PCR reagents and PCR cycling conditions.

**Table 1.** Antibodies and the dilutions used for flow cytometry.

| Specificity | Clone | Fluorochrome | Dilution | Volume per sample (µl) |
| --- | --- | --- | --- | --- |
| CD3 | UCHT1 | Alexa Fluor 488 | 1:200 | 0.25 |
| CD62L | DREG-56 | PerCP-Cy5.5 | 1:50 | 1 |
| CD19 | HIB19 | Pacific Blue | 1:200 | 0.25 |
| CD14 | M5E2 | Pacific Blue | 1:200 | 0.25 |
| CD56 | MEM-188 | Pacific Blue | 1:200 | 0.25 |
| CD11c | B-ly6 | V450 | 1:200 | 0.25 |
| CD8a | RPA-T8 | Brilliant Violet 570 | 1:100 | 0.5 |
| PD-1 | EH12.1 | PE-CF594 | 1:25 | 2 |
| CD45RA | HI100 | PE-Cy7 | 1:50 | 1 |
| TIM-3 | 7D3 | Alexa Fluor 647 | 1:25 | 2 |
| CD4 | SK3 | APC-H7 | 1:100 | 0.5 |

### 12. Flow cytometry profiling of cells from the platform (day 8 or as preferred)

#### Timing: [3h]

This step describes the process of sample preparation and staining for flow cytometry to study and characterize T cell populations present in the edited cells at the end of the pipeline.

1. Centrifuge cells at 400 g for 5 min at 4 °C.
  a. **Note:** we recommend 0.5-1 million cells/sample for flow staining.
2. Discard supernatant and resuspend cells in 200 µl cold PBS.
3. Transfer cell suspension into a 96-well V bottom plate.
4. Centrifuge cells at 400g for 5 min at 4 °C.
5. While centrifuging, prepare 50 µl of 1/500 dilution of Live/Dead Violet dye in PBS with 1/20 dilution of human Fc block for each sample (e.g., 2 µl Live/Dead + 50 µl Fc block per 1000 µl PBS for 20 samples).
  a. **Note:** Do not re-freeze the thawed stock dye solution, discard any leftover.
6. Discard the supernatant.
7. Add 50 µl of the Live/Dead Dye + Fc block mix to each sample. Mix well. Stain for 30 min 4° C on ice & dark.
8. After the 30 min of live/dead staining + Fc block, add 100 µl of PBS to each sample.
9. Centrifuge at 400 g for 5 min at 4°C.
10. While centrifuging, prepare the antibody mix (Table 2). Add 41.75 µl of the flow staining buffer to the antibody mix to achieve staining volume of 50 µl.
11. Discard the supernatant and add 50 µl of antibodies mix to each sample. Mix well.
12. Stain for 30 min 4°C on ice & dark.
13. While staining cells, prepare compensation controls:
14. Prepare compensation controls:
  a. Completely resuspend the AbC™ Total Compensation capture beads (Component A) and negative beads (Component B) by gently vortexing for 10 s before use.
  b. Label flow tubes for each fluorochrome and additional tube for unstained control.
  c. Add one drop of AbC™ Total Compensation capture beads (Component A) to each tube.
  d. Add 1 µl of each antibody conjugate to the AbC™ Total Compensation capture bead suspension in the designated tube and mix well. Make sure to deposit the antibody directly to the bead suspension. Incubate for 15 min at RT, protected from light.
  e. Add 3 mL of PBS to sample tubes. Centrifuge for 5 min at 250 g.
  f. Carefully remove the supernatant from tubes and resuspend the bead pellet by adding 0.5 mL of PBS to sample tubes.
  g. Add one drop of negative beads (Component B) to each of the tubes and mix well.
  h. Vortex tubes before analysing using flow cytometry to increase percentage of singlet beads.
15. After the antibody staining, add 150 µl of cold flow staining buffer to each sample.
16. Centrifuge at 400 g for 5 min 4°C.
17. Discard supernatant.
18. Resuspend cells in 200 µl of cold flow staining buffer.
19. Transfer samples into flow tubes.
20. Analyze the samples in LSRII (BD Bioscience) or other flow cytometry instrument.

### 13. Cryopreservation of corrected cells

#### Timing: [30 min]

This is the final step to the protocol, where edited cells from the platform are cryopreserved for long-term storage. We recommend cryopreserving a minimum of 1 million cells/vial (e.g. cells used for flow cytometry) and a maximum of 30 million cells/vial (e.g. cells thawed for further in vitro experimentation) for optimal cell recovery.

1. Follow cryopreservation steps as described previously for PBMC cryopreservation in Step 3.
  a. **Note:** We recommend following the guidelines below when thawing the cells for further assessments:
    i. Flow cytometry: Thaw cells as described in Step 4 at 1 million cells/mL in ImmunoCult™-XF T Cell Expansion Medium supplemented with 250 U/mL IL-2 and allow to rest o/n at 37 °C/5 % CO_2_ before proceeding with cell staining and flow cytometry.
    ii. Re-stimulation for further experimentation: Thaw cells as as described in Step 4 at 1 million cells/mL in T cell stimulation medium and allow to rest for 2-3 days before further expansion or intended experiment.

#### Expected outcomes

Following the instructions of this protocol, we expect that by the end of the platform (day 8 and beyond) CD3+ T cells account for ≥ 80% of all cells, where frequencies of CD4+ and CD8+ T cells depend on the individual [1]. Under the PBMC culturing conditions, we expect cells to proliferate in a way that starting with very low PBMC material (1-5 million) will result in a robust proliferation and optimal viability (Fig. 2). Furthermore, under these culture conditions, we also expect to observe T cells with low exhaustion levels and that central and effector memory T cells persist in the culture (Fig. 3-4). Finally, we expect to see up until 50% HDR in the edited T cells with optimized CRISPR/Cas9 reagent design, and up to 80% when NHEJ inhibitors are used [1]. However, we note that when working with patient cells with different pathogenic mutations, individual variation in terms of T cell proliferation, viability, exhaustion/ phenotype and editing should be expected.

**Figure 3.**
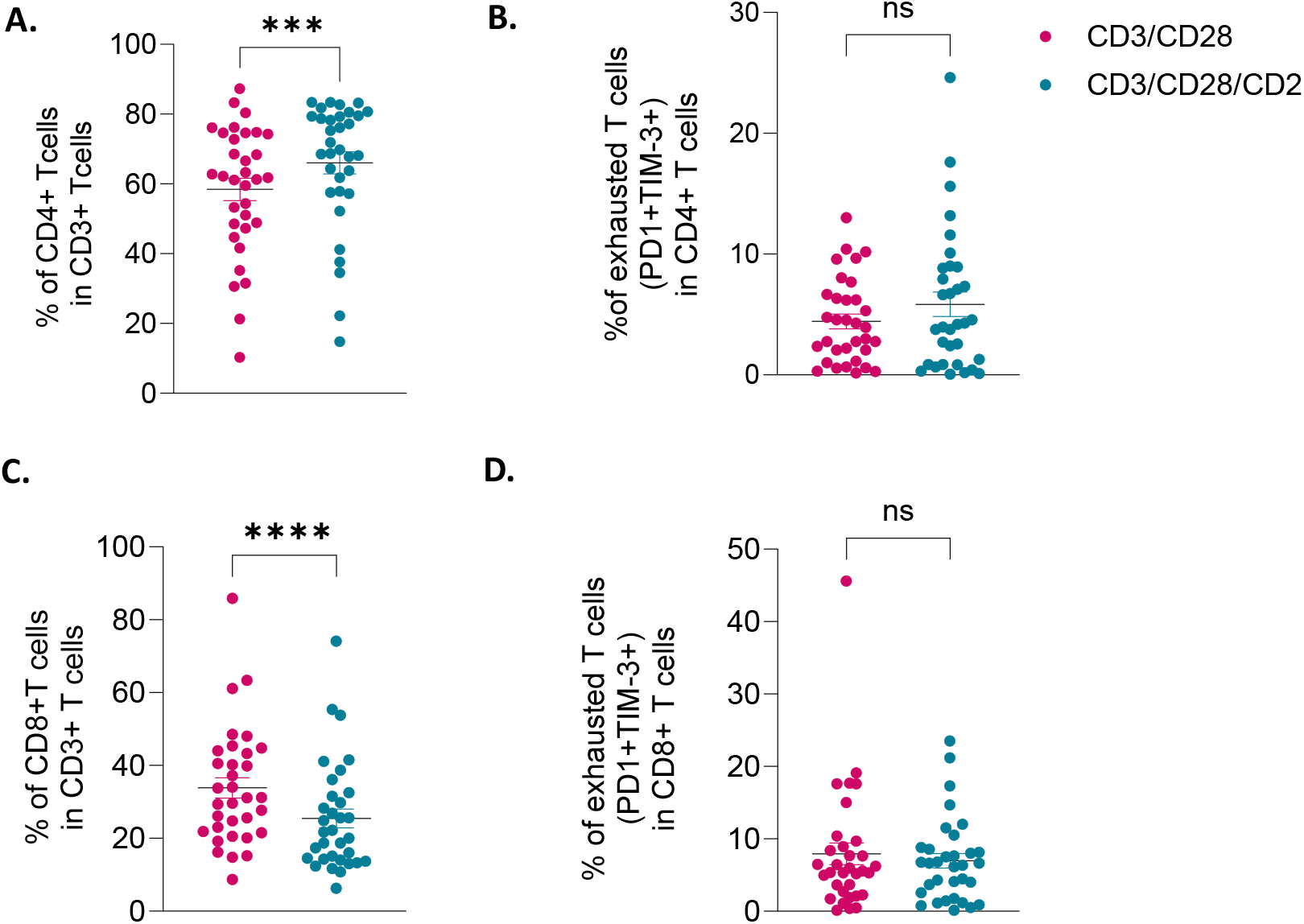
Assessment of T cell exhaustion. Cells generated from cells from 33 IEI patients as described in figure 2A were cryopreserved on day 8 of the PBMC stimulation and cell culture. The cells were thawed and analyzed by flow cytometry. (A) CD4^+^ cells among CD3^+^ T cells. (B) Exhausted (PD-1^+^TIM-3^+^) cells among CD4^+^ T cells. (C) CD8^+^ cells among CD3^+^ T cells. ( (D) Exhausted (PD-1^+^TIM-3^+^) cells among CD8^+^ T cells. Data are presented as mean ± SEM. Statistical analyses were performed in GraphPad Prism using the Wilcoxon matched-pairs signed rank test (nonparametric, paired design). P values were adjusted for multiple comparisons using the Holm–Šidák method, with a significance threshold of α = 0.05.

**Figure 4.**
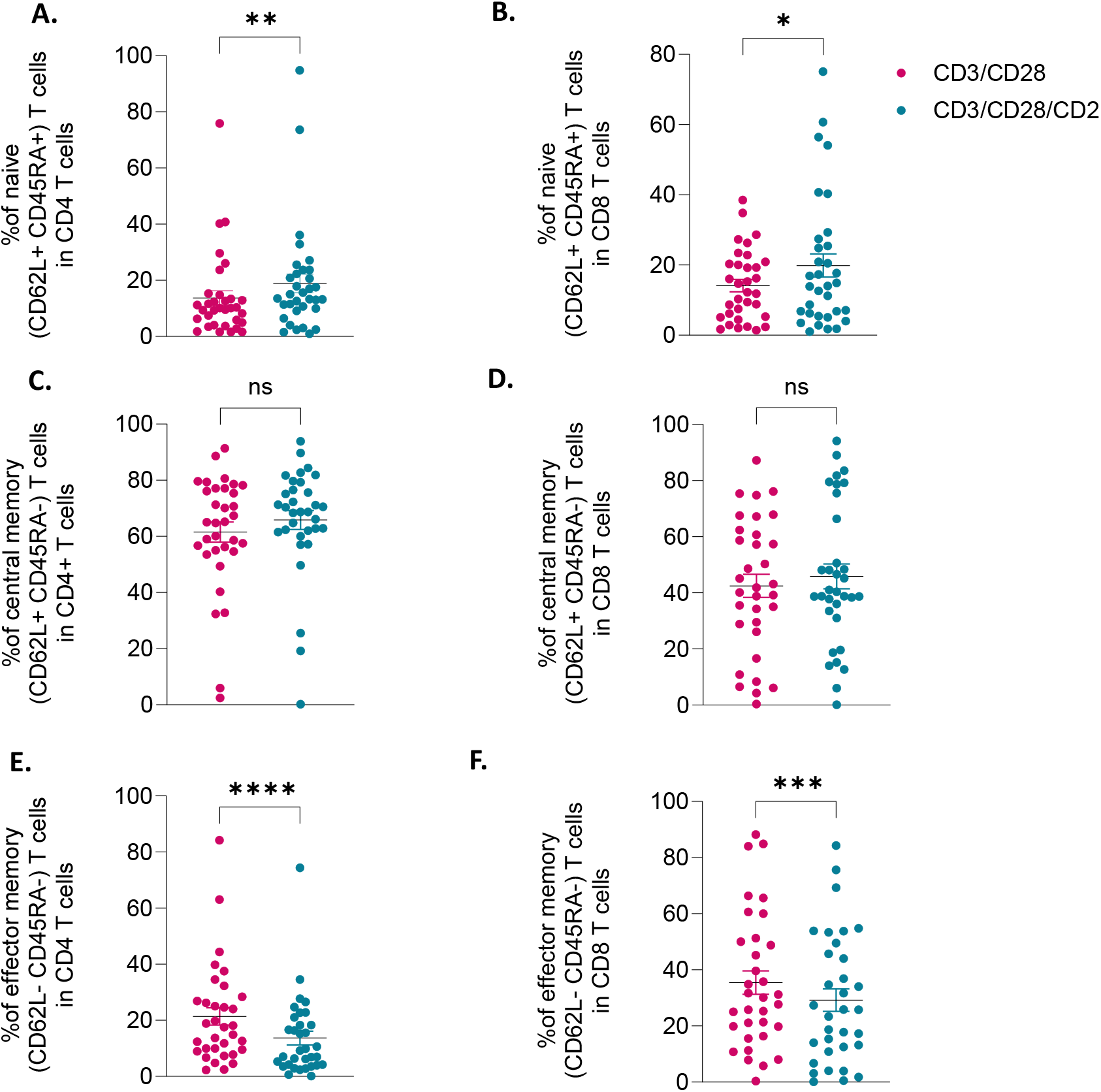
Assessment of phenotype of T cells. Cells generated from cells from 33 IEI patients as described in figure 2A were cryopreserved on day 8 of the PBMC stimulation and cell culture. The cells were thawed and analyzed by flow cytometry. (A) Naïve (CD62L^+^CD45RA^+^) cells among CD4^+^ T cells (B) Naïve (CD62L^+^CD45RA^+^) cells among CD8^+^ T cells (C) Central memory (CD62L^+^CD45RA^−^) cells among CD4^+^ T cells (D) Central memory (CD62L^+^CD45RA^−^) cells among CD4^+^ T cells (E) Effector memory (CD62L^-^CD45RA^−^) cells among CD4^+^ T cells and (F) Effector memory (CD62L^-^CD45RA^−^) cells among CD4^+^ T cells. Data are presented as mean ± SEM. Statistical analyses were performed in GraphPad Prism using the Wilcoxon matched-pairs signed rank test (nonparametric, paired design). P values were adjusted for multiple comparisons using the Holm–Šidák method, with a significance threshold of α = 0.05.

#### Key resources table

**Key resources table:**
| Reagent or resource | Source | Identifier |
| --- | --- | --- |
| Lymphoprep | Stemcell Technologies | #18061 |
| LeukoSep tube (50 mL) | Greiner | #227288 |
| PBS (pH 7.4) | Thermo Fisher Scientific | #10010023 |
| 40 μm Cell Strainer | Corning | #352340 |
| CryoStor freezing medium | Stemcell Technologies | #100-1061 |
| ImmunoCult™-XF T Cell Expansion Medium | Stemcell Technologies | #10981 |
| ImmunoCult™ Human CD3/CD28 T Cell Activator | Stemcell Technologies | #10971 |
| ImmunoCult™ Human CD3/CD28/CD2 T Cell Activator | Stemcell Technologies | #10970 |
| Recombinant human IL-2 | Miltenyi Biotec | #130-097-742 |
| Recombinant human IL-7 | Miltenyi Biotec | #130-093-937 |
| Recombinant human IL-15 | Miltenyi Biotec | #130-093-955 |
| Alt-R S.p. Cas9 Nuclease V3 | Integrated DNA Technologies | #1081058 |
| Armadillo PCR plate (96-well) | Thermo Fisher Scientific | #AB3396 |
| Microplate Aluminum Sealing Tape | Corning | #6570 |
| TheraPEAK™ P3 solution set | Lonza | #G4LP3-22500 |
| DNeasy Blood & Tissue Kit | Qiagen | #69506 |
| Qubit™ dsDNA HS Assay Kit | Qiagen | #Q32854 |
| ddPCR plates, 96 well, semi-skirted | Bio-Rad | #12001925 |
| ddPCR Supermix for Probes (No dUTP) | Bio-Rad | #1863025 |
| Piercable Foil Heat Seal | Bio-Rad | #1814040 |
| Automated Droplet Generator | Bio-Rad | N/A |
| PX1 PCR Plate Sealer | Bio-Rad | N/A |
| QX200™ Droplet Reader | Bio-Rad | N/A |
| QX Manager | Bio-Rad | N/A |
| 96-well V bottom plate | Corning | CLS3956 |
| CoolCell container | Corning | N/A |
| 15 mL Falcon tube | Corning | #352096 |
| 50 mL Falcon tube | Corning | #10788561 |
| SafeSeal reaction tube, 1.5 mL | Sarstedt | #72.706.006 |
| Lonza 4D-Nucleofector | Lonza | N/A |
| Lonza 16-well strip or 96-well nucleofection plate | Lonza | N/A |
| TheraPEAK™ P3 solution set | Lonza | #G4LP3-22500 |
| AbC™ Total Antibody Compensation Bead Kit | Invitrogen | #A10513 |
| LIVE/DEAD™ Fixable Violet Dead Cell Stain Kit, for 405 nm excitation | Thermo Fisher | #L34963 |
| Alexa Fluor® 488 anti-human CD3 Antibody | Biolegend | #300415 |
| PerCP/Cyanine5.5 anti-human CD62L Antibody | Biolegend | #3004823 |
| Pacific Blue™ anti-human CD19 Antibody | Biolegend | #302232 |
| Pacific Blue™ anti-human CD14 Antibody | Biolegend | 301828 |
| Pacific Blue™ anti-human CD56 (NCAM) Antibody | Biolegend | 304629 |
| Brilliant Violet 570™ anti-human CD8a Antibody | Biolegend | 301037 |
| PE/Cyanine7 anti-human CD45RA Antibody | Biolegend | 304125 |
| BD Horizon™ V450 Mouse Anti-Human CD11c | BD Biosciences | 560369 |
| BD Horizon™ PE-CF594 Mouse Anti-Human CD279 (PD-1) | BD Biosciences | 565024 |
| BD Pharmingen™ Alexa Fluor® 647 Mouse Anti-Human TIM-3 (CD366) | BD Biosciences | 565559 |
| BD™ APC-H7 Mouse Anti-Human CD4 | BD Biosciences | 641398 |
| FcR Blocking Reagent human | Miltenyi | 130-059-901 |
| eBioscience Flow Cytometry Staining Buffer | Invitrogen | #00-4222-26 |
| 5 mL Round Bottom Polystyrene Test Tube | Corning | #352054 |
| LSR II Flow Cytometer | BD Biosciences | N/A |

#### Limitations

There are certain limitations to the study, which are discussed here. First, regarding CRISPR gRNA design using SpCas9, PAM availability determines the number of possible gRNAs to test per given locus. If SpCas9 PAM sites are not available, please consider using another source of Cas9 (e.g. SaCas9), engineered Cas9 with wider PAM availability (e.g. SpRY-Cas9) or another Cas variant (e.g. Cas12a). Second, it is not possible to integrate synonymous SNVs in the repair template strategy for noncoding mutations. This can result in reduced on-target HDR and potentially issues in accurate detection by ddPCR. To mitigate this, we recommend considering alternative HDR-based assays (e.g. amplicon sequencing) and using other HDR enhancing strategies, such as NHEJ inhibition. However, we advise to assess potential safety issues (e.g. cell viability, chromosome arm loss etc.) on a case-by-case basis when using NHEJ inhibitors. Third, there may be individual variation in T cell response to cytokine stimulation presented in the protocol. We expect mutations that affect cytokine signalling (IL-2, IL-7, IL-15) and co-stimulation (CD3/CD28 or CD3/CD28/CS2) to require an alternative and user-defined stimulation setup to promote cell proliferation.

#### Troubleshooting

**Problem 1:** PBMCs are aggregated and difficult to pipette into single-cell suspension for counting and cryopreservation.

**Solution:** Strain cells through a 40 μM cell strainer to ensure cells are in single cell suspension before counting and cryopreservation to remove the aggregates.

**Problem 2:** A high amount of red blood cells are present in the PBMCs after density grade centrifugation.

**Solution:** A red blood cell lysis buffer can be used to eliminate contaminating red blood cells at this stage. Use the reagent of choice and follow the vendor’s guidelines. If the red blood cells are present but they don’t significantly affect the cell pellet size, the lysis step is not necessary.

**Problem 3:** PBMC viability is poor (<70%) on Day 4-8 after thawing despite good cell viability upon thawing (>90%).

**Solution:** If cell viability is drastically decreased upon cell culture and persist by day 8 of the platform, check if the mutation alters cytokine signalling and the subsequent responsiveness to cytokines used in this protocol, as this could result in reduced cell viability and proliferation. When necessary, find alternative T cell stimulation cocktail to promote T cell proliferation from PBMCs. If mutation does not point to abnormal responses to cytokines, cytokines used in this protocol can be titrated to preserve cell viability but this can result in suboptimal cell proliferation.

**Problem 4:** There are too few cells to nucleofect on day 4 of the T cell platform.

**Solution:** If there are not enough cells for nucleofection on day 4 of the cytokine stimulation, dilute cells on day 4 as instructed in Step 5 and repeat cell dilution again on day 6 if necessary. Alternatively, as low as 0.1 Million cells can be used for nuclefection, as described in Step 6.

**Problem 5:** HDR editing is low.

**Solution:** Low HDR editing can result from several factors, including low cell viability prior to and after nucleofection and competition between HDR and NHEJ upon DNA repair. If cell viability is low (<70%) upon nucleofection, cells typically will not survive well after nucleofection. Consider trying a different nucleofection code for T cells (consult nucleofector vendor), a different nucleofection instrument (e.g. MaxCyte) or an alternative nucleofection buffer. If cell viability is good after editing, it is possible that NHEJ editing is dominant and resulting HDR is low. To mitigate this, NHEJ inhibitors, such as KU0060648 at 0.5 μM 24h after nucleofection [1] . As NHEJ also inhibit cell proliferation, remove compound 24h after nucleofection by changing the medium or remove half of the medium and replace it by fresh T cell recovery medium to dilute the compound in culture.

**Problem 6:** NHEJ editing is low when the desired editing outcome is a knockout.

**Solution:** After gRNA validation for the gRNA sequence with the highest NHEJ efficiency, it is possible that edited cells have a selective disadvantage in cell culture upon editing. In some of those cases, it is possible that the cells with the knockout disappear in culture over time and thus, we advise performing a time course experiment for all new loci to assess the optimal timepoint for collecting the edited cells.

## Supporting information

Graphical abstract

## Resource availability

### Lead contact

Further information and requests for resources and reagents should be directed to and will be fulfilled by the lead contact, Emma Haapaniemi.

### Technical contact

Technical questions on executing this protocol should be directed to and will be answered by the technical contact, Shiva Dahal-Koirala.

## Materials availability

All reagents used in this study are available from the lead contact with a completed materials transfer agreement.

## Data and code availability

This protocol does not report any original code.

## Acknowledgements

We thank all patients and families for their participation in the study. We thank Karolinska Institute Protein Science Facility for manufacturing the Cas9 protein, and Riitta Lehtinen for her expert technical assistance. Research Council of Norway, Health South-East Region, Swedish Childhood Cancer Society (Barncancerfonden), and Norwegian Cancer Society supported this work. This work was partially supported by the Research Council of Norway through its Centres of Excellence scheme (project number 332727).

## Author contributions

**K.M**. performed and analysed most of the experiments that established the T cell platform and wrote the manuscript. **A.S.M** performed T cell culture experiments in IEI patients. **S.D.K** designed flow cytometry experiments, performed, analysed & supervised research and wrote the manuscript. **E.H**. generated funding and supervised the study. All authors read and approved the manuscript.

