## Supplementary figures and images for "Protocol for Non-viral HDR-based CRISPR/Cas9 platform for small custom editing in primary T cells"

### Graphical abstract

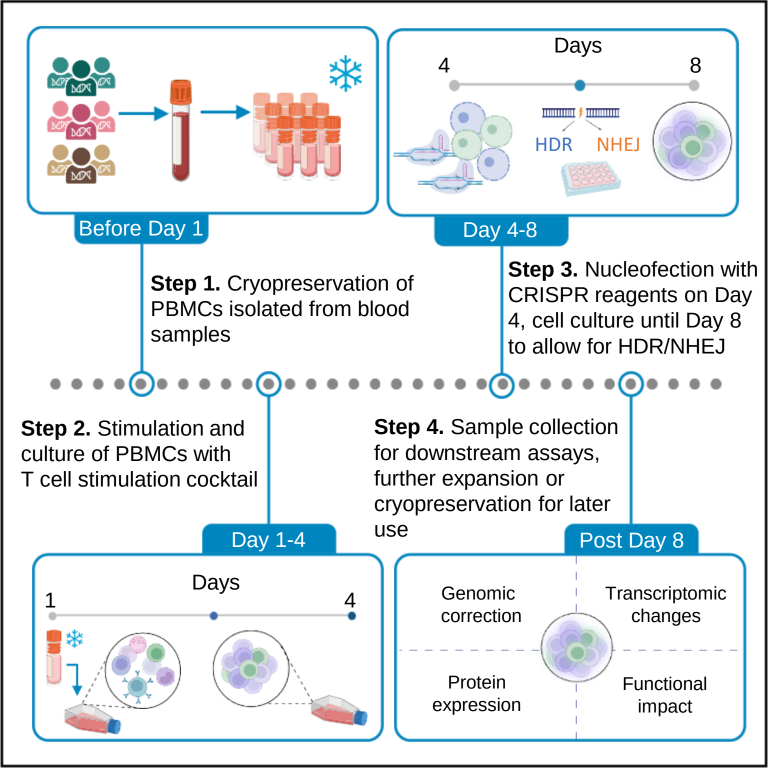
